# *SigRescueR*: A Pan-System Framework for Noise Correction and Mutational Signature Identification Across Sequencing Platforms

**DOI:** 10.1101/2025.11.14.688578

**Authors:** Peter T. Nguyen, Maria Zhivagui

**Author notes:** Corresponding author: Maria Zhivagui.

## Abstract

Mutational signatures serve as molecular fingerprints of the biological processes and exposures that shape cancer genomes. However, accurate signal recovery remains challenging due to pervasive background variants, sequencing artifacts, technical noise, and platform-specific biases that obscure true mutagenic patterns, hampering biomarker discovery and mechanistic interpretation. Here we introduce *SigRescueR*, a rigorous, pan-system, computational framework designed for noise correction and mutational signature identification. *SigRescueR* applies statistically robust baseline correction to effectively disentangle true mutational signals from confounding noise and artifacts. When applied to extensive datasets spanning experimental models and human cancers, *SigRescueR* reliably identified canonical mutational signatures associated with environmental mutagens such as colibactin, benzo[a]pyrene, and UV radiation, and chemotherapeutic agents, namely 5-fluorouracil and cisplatin. *SigRescueR* effectively operated across diverse mutation classes, including single base substitutions, insertions and deletions, and doublet base substitutions, while also integrating strand bias and duplex sequencing data for toxicology applications. *SigRescueR* offers a unified, high-precision platform that seamlessly integrates cancer genomics, molecular toxicology, and mechanistic studies. It enables precise mapping of mutagenic processes and identification of robust genomic biomarkers of environmental and therapeutic exposures, providing a transformative framework for translational cancer research.

**Availability and implementation:** *SigRescueR* is implemented in R and provided as open-source software on GitHub at https://github.com/ZhivaguiLab/SigRescueR/

## Introduction

Cancer is fundamentally driven by somatic mutations arising as the result of Darwinian evolution where a single cell evades growth-control mechanisms and begins to divide uncontrollably [1–4]. These somatic mutations result from mutational processes that are characterized by an interplay of DNA damage, repair, and replication [5–9]. The origins of these mutational processes are diverse, encompassing both exogenous factors, such as environmental carcinogens, and endogenous sources including cellular aging and genomic instability [10–14]. Advances in analyzing mutational patterns in cancer genomes have revealed the sources of somatic mutations and identified key risk factors [5,10,15–20]. These signatures are systematically cataloged in the COSMIC database, serving as a key resource for cancer genome studies [10]. Experimental studies, both *in vivo* and *in vitro*, have validated several signatures observed in cancer patients, and uncovered novel mutagenic exposures and mechanisms [13,14,21–27]. However, a persistent challenge is that these mutational signatures often contain substantial noise arising from endogenous processes [7,28–30], cell culture artifacts [31], and sequencing errors [32]. This confounding background noise introduces variability between samples, distorting the true biological mutational signatures. Moreover, extensive analysis of 23,829 samples (19,184 whole-exome and 4,645 whole-genome sequencing) have identified 19 single-base substitutions (SBS) artifactual signatures [10], including batch effects (SBS95) [33], germline variant associations (SBS54) [10], and sequencing artifacts (SBS45) [10,32]. Therefore, effective removal of baseline and artifactual mutations is essential not only for experimental models but also for tumor samples, to reveal subtle genomic patterns and uncover hidden insights into cancer biology.

To address this challenge, two strategies have been employed to remove baseline mutational processes. The first relies on straightforward exposure-baseline subtraction, which assumes that mutations absent in the baseline are exposure-associated. However, this method falters when the baseline mutation burden exceeds the exposure, producing negative values that must be replaced by zeros [25,34]. The second strategy leverages non-negative matrix factorization (NMF) [35] to decompose mutation matrices and separate control from exposure-related signatures [36]. Yet, NMF’s performance depends heavily on sample size [37], rendering it impractical with limited replicates [38]. Another approach that can be considered is the non-negative least squares (NNLS) algorithm [39]. NNLS can assign background activity which can then be subtracted from exposed samples. This method is particularly well-suited for analyses of individual samples. Nevertheless, NNLS’s propensity to overfit can lead to negative values [40]. Collectively, these limitations highlight the urgent need for a burden-insensitive approach that can effectively remove baseline effects without producing negative attributions, even when the background profile dominates or closely resembles the treatment profile.

We developed *SigRescueR*, a robust R package that applies Bayesian inference to remove baseline mutational patterns from exposure-derived mutational profiles. We demonstrate that *SigRescueR* preserves the integrity of the original data, enabling near-perfect reconstruction of mutational profiles, including accurate retention of mutational burden. By leveraging COSMIC signatures as a reference standard, *SigRescueR* significantly improved similarity of the inferred treatment signatures to known ground truths. We demonstrate that *SigRescueR* supports a wide range of SBS classifications, including SBS96 and SBS288 channels, demonstrating impressive versatility. In rigorous comparisons, *SigRescueR* outperformed existing tools while uniquely providing credible intervals that quantify uncertainty, a crucial advantage over methods that offer only point estimates. *SigRescueR* adapts seamlessly across diverse models, species, patients’ data, and sequencing platforms, including cutting-edge duplex DNA sequencing, making it an indispensable asset for mutational signature analysis and toxicology studies.

## Method

### Overview of *SigRescueR*

*SigRescueR* was developed to disentangle baseline mutational patterns from those induced by exposure or treatment (**Figure 1**). The algorithm employs Bayesian inference to jointly model both the reconstructed mutational profile and the total mutation count at the mutational context level. Specifically, *SigRescueR* utilizes count-based observed mutational spectra (“Observed”) and a proportion-based background mutational signature (*s_b_*). The reconstructed mutational profile (“Reconstructed”) is represented as a combination of the background signature *s_b_*and latent exposure signature (*s_e_*), each weighted by their respective activity parameters (θ*_b_*, θ*_e_*), and scaled by the observed mutation burden to recover mutation counts.

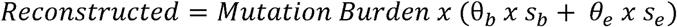

**Figure 1.**
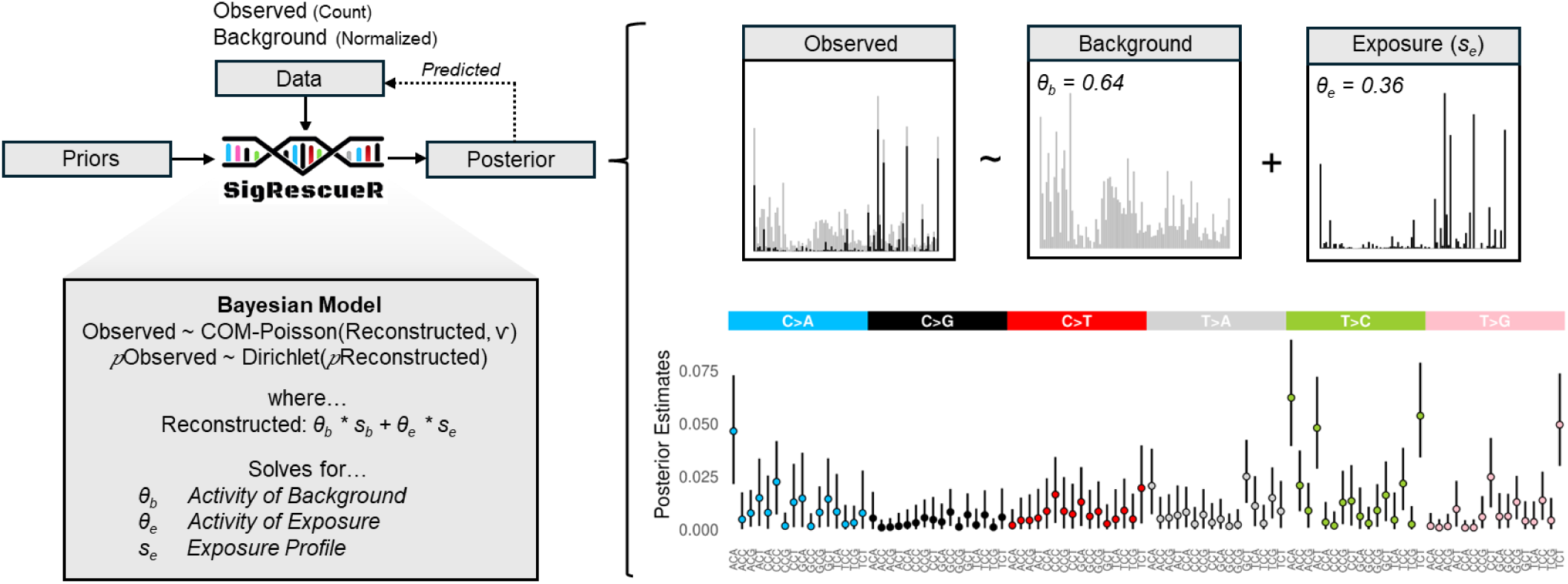
Overview of *SigRescueR*. Using the observed (exposed sample) and background (unexposed sample or provided) mutational profiles, *SigRescueR* applies Bayesian inference with multiple iterations and chains to subtract the background profile (grey bars) from the observed profile (a mixture of exposure- and endogenous-associated mutations). This process rescues the exposure-associated signal (black bars). The method utilizes count data from the exposed sample and the normalized background profile to estimate both background and exposure activities while inferring the exposure-associated profile. By exploring the parameter space, *SigRescueR* generates a full posterior distribution capturing uncertainty inherent in the data.

*SigRescueR* incorporates a Gamma prior that assumes θ*_b_* is higher than θ*_e_*. This prior enforces positivity and models a skewed distribution centered around 1 for θ*_b_* and 0.2 for θ*_e_*. This design prevents the exposure signature from dominating the reconstructed profile by absorbing the majority of the signal.

The observed mutation spectra proportions were modeled using a Dirichlet distribution parameterized by the reconstructed mutation spectra proportions.

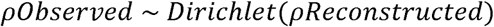

The observed counts were modeled using a Conway-Maxwell Poisson (COM-Poisson) distribution [41] parameterized by the reconstructed mutation counts. The dispersion parameter (□) of the COM-Poisson was constrained with a Normal prior centered at 2, with a lower bound of 1.1 to capture underdispersion and to penalize large deviations from the observed counts.

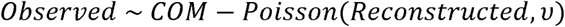

To encourage accurate reconstruction, parameter configurations yielding a cosine similarity greater than 0.95 between the reconstructed and observed profiles were positively rewarded.

To accommodate variability among samples with the same exposure, a weighted mutational profile was generated where each sample (*i)* contributed proportionally to its total mutation burden (TMB). The weighted profile was computed as the sum of each sample’s mutational profile (*mi*) multiplied by its corresponding weight (*wi*).

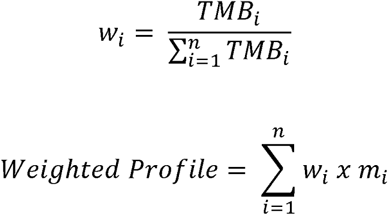

### *SigRescueR* implementation

*SigRescueR* is implemented in R [42] and applies Bayesian inference using Rstan [43]. Bayesian inference is configured to run four Markov Chain Monte Carlo (MCMC) chains with 2000 warm up iterations followed by 4000 sampling iterations for each chain, which can be reconfigured within the SigRescueSetup function. To reduce runtime, *SigRescueR* supports parallelizing computation of up to 20 samples implemented in parallel [42]. The SigRescueAnalyze function processes the output of SigRescueRun function by extracting the median of θ*_b_* and θ*_e_* and the lower limit (2.5% credible interval) of *s_e_* as representative estimates.

### Evaluation of reconstruction

To evaluate reconstruction accuracy, we benchmarked the reconstructed against the original spectra using count accuracy, Jensen-Shannon divergence (JSD) [44], and cosine similarity [45]. To assess the recovery of true biological signals, similarity between the exposure signature and COSMIC signatures was performed. JSD quantifies differences in relative proportions, where values near 0 denote high similarity.

### Benchmarking cleaning performance

To assess *SigRescueR’s* performance, we provided identical input for direct comparison against simple subtraction [25,34], NNLS [46], and SparseSignatures [47] (**Supplementary Table 1**). The simple subtraction method subtracts the baseline mutational spectra from the exposed mutational spectra and clips negative values resulting from the subtraction to zero, representing the most basic form of cleaning. NNLS is an optimization method constrained to estimate the baseline activity under non-negative constraint while minimizing the least squared error. SparseSignatures employs an NMF framework, incorporating a *LASSO* penalty, for mutational signature discovery based on a predefined number of signatures while also estimating the activity for each signature. Count accuracy, reconstruction cosine similarity, and COSMIC cosine similarity were used as metrics to evaluate performance. Count accuracy was measured as the difference in mutation counts where values close to 0 indicate minimal deviation from the original mutation count. Reconstruction cosine similarity was computed between the reconstructed and original mutational spectra. COSMIC cosine similarity was computed between the inferred exposure mutational signature and COSMIC signature.

### Synthetic injection of SBS signatures

We randomly selected 100 samples confirmed to be negative for both SBS45 and SBS54 signatures defined by SigProfilerAssignment [48]. For each sample, the level of synthetic injection ranged between 5%-95%, representing the fraction of the total mutational profile composed of synthetic SBS45 or SBS54.

After injection, we evaluated *SigRescueR*’s ability to identify injections while preserving the original spectra as measured by the precision (accuracy of true positives) and sensitivity (detection of true positives). We also computed the F1 score to assess the balance between precision and sensitivity.

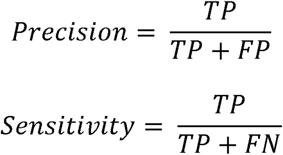

where TP denotes correctly identified injections, FP denotes misclassified injections, and FN denotes missed injections.

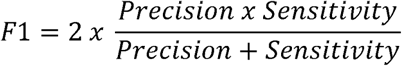

### Statistics and reproducibility

All pairwise statistical comparisons were performed using Mann–Whitney U test [49]. Overall statistical differences between groups were performed using Anova [50]. JSD and cosine similarity were computed to quantify the similarity. All SBS attributions were performed using SigProfilerAssignment [48].

## Result

### Baseline correction increases similarity to reference

*SigRescueR* implements statistically rigorous baseline correction to separate true mutational signatures from background noise and technical artifacts, significantly improving the detection of biologically meaningful mutational processes (**Figure 1**). To evaluate how well *SigRescueR* removes baseline mutations, we analyzed diverse experimental models, including induced pluripotent stem cells (iPSCs) [51], immortalized N/TERT keratinocytes (NTERT1) [52], human intestinal organoids (HIO) [25], mouse embryonic fibroblasts (MEFs) [21], and mouse intestinal organoid [53] (**Supplementary Table 2**). These models represent various exposures with known COSMIC signatures (**Supplementary Table 3**). The samples exhibited highly variable mutational burdens, ranging from 369 to 4,427 mutations in human models (n = 26) and from 595 to 24,947 mutations in mouse models (n = 23). For each model, samples treated with non-mutagenic conditions were used as backgrounds. We, first, used colibactin-treated HIO to extract the associated mutational signature and reconstruct the original profile (**Figure 2A**). The reconstructed profile demonstrated close agreement with the observed profile with a cosine similarity of 0.963. The posterior mean activities were 0.474 for exposure and 0.526 for background signatures, supported by convergence across chains (**Supplementary Figure 1A-C**).

**Figure 2.**
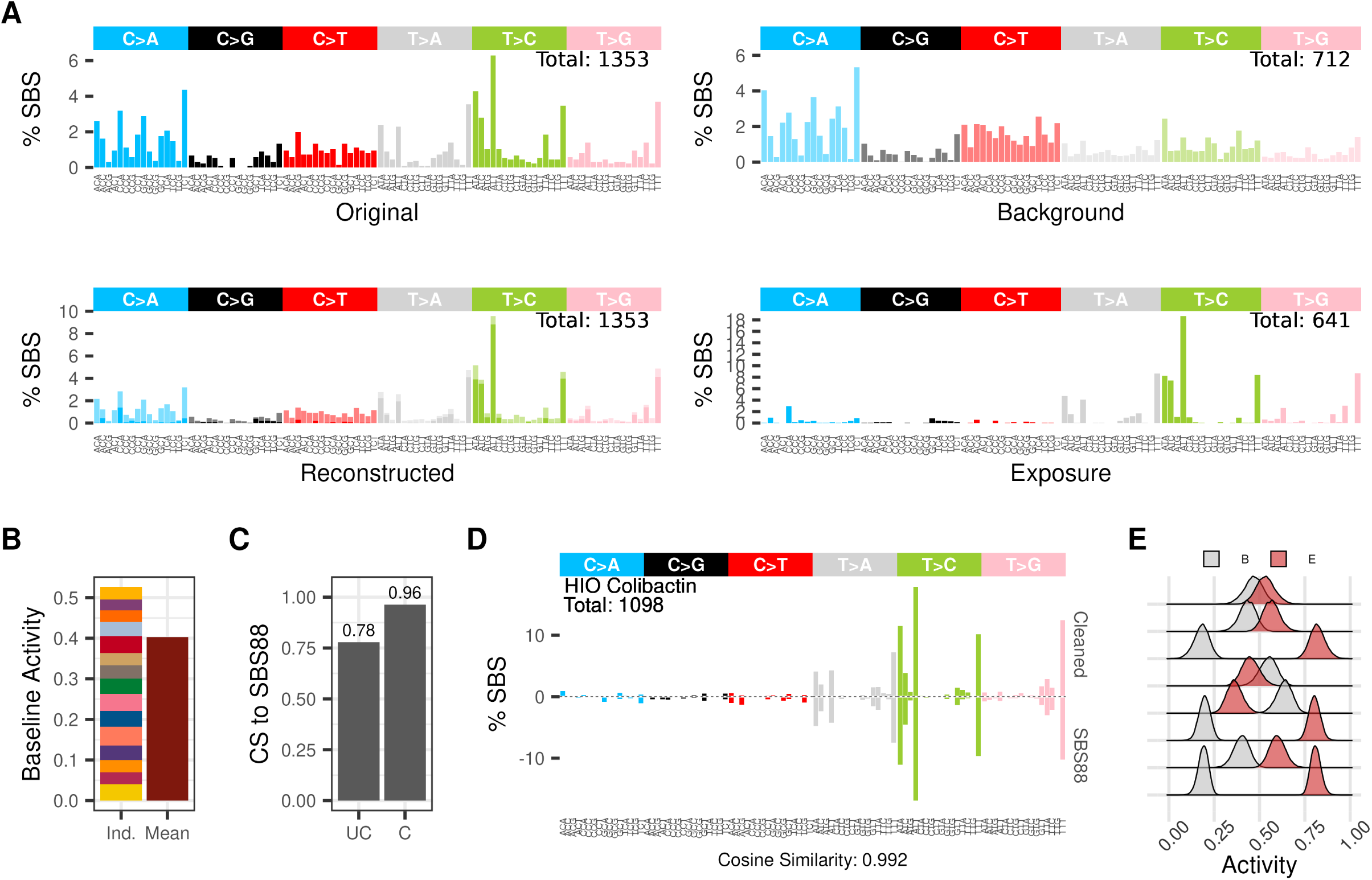
Background cleaning of a human intestinal organoid sample exposed to colibactin. **(A)** SBS-96 mutational signature plot showing the uncleaned (Original), control (Background), inferred treatment (Exposure), and reconstructed profiles. Alpha (⍰) transparency indicates background (⍰ = 0.5) and exposure (⍰ = 1) levels in the reconstructed profile. **(B)** Barplot depicting relative background activity estimated using multiple backgrounds (Ind.) and a single background (Mean), with fill color denoting individual replicates in the Ind. column. **(C)** Barplot showing cosine similarity to the reference signature (SBS88) before (UC = uncleaned) and after baseline correction (C = cleaned). **(D)** Weighted SBS-96 mutational signature plot comparing the inferred exposure (Cleaned) with the COSMIC reference (SBS88). The total mutation count in the cleaned profile is displayed within the plot. Cosine similarity scores reflect similarity between the cleaned signature and SBS88. **(E)** Posterior distributions of exposure (red-filled densities: E) and background activity (grey-filled densities: B) across human intestinal organoid replicates, with each ridge representing a single sample.

Moreover, baseline correction was further evaluated by comparing the use of multiple background profiles to a single composite background profile, generated by a weighted combination of all available background profiles. Results indicated that baseline correction utilizing multiple profiles yielded a higher background activity parameter (θ*_b_* = 0.526) compared to correction using a single profile (θ*_b_* = 0.402) (**Figure 2B**). As expected, this enhancement is likely attributable to the greater ability of multiple backgrounds to capture variability in the observed mutational profiles that a single background may not represent (**Supplementary Figure 2**). To assess consistency, *SigRescueR* was applied to the same sample across 100 independent iterations. The residuals of exposure activity were consistently minimal (residual < 0.0011), and the inferred exposure mutational signature exhibited high stability (cosine similarity > 0.99) (**Supplementary Figure 1D-E**), substantiating the robustness of the method. To evaluate the effectiveness of baseline correction, we computed the cosine similarity between SBS88, a colibactin-induced COSMIC mutational signature [25,26,54], and the colibactin-exposure signature inferred by *SigRescueR*. Baseline correction substantially improved the cosine similarity from 0.78 to 0.96, a 23% enhancement (**Figure 2C**). Further, when aggregating the weighted mutational profiles from all colibactin-exposed HIO (n = 8), the resulting profile exhibited a cosine similarity of 0.992 to SBS88, indicating near-perfect concordance (**Figure 2D**). Notably, assessment of exposure activity across biological replicates demonstrated pronounced variability, with several replicates showing higher background activity than exposure-related activity, highlighting biological and technical heterogeneity within experimental replicates from a single study (**Figure 2E, Supplementary Figure 3A**).

### Baseline correction achieves high reconstruction fidelity

Following the strong concordance observed between the inferred exposure signature and the COSMIC reference, reconstruction accuracy was systematically assessed by comparing mutation counts and profile similarity between the reconstructed and observed mutational spectra. Our analysis revealed a strong linear correlation in mutation counts across samples (**Figure 3A**). Furthermore, *SigRescueR* achieved a near-perfect cosine similarity of 0.99 between observed and reconstructed mutational profiles (**Figure 3A**), confirming the high fidelity of the reconstruction. To evaluate reproducibility, we examined concordance of the inferred exposure mutational signatures across models and replicates. Consistent high cosine similarity among replicates within each model reflected robust reproducibility (**Figure 3B**). Notably, exposure signatures generated from distinct models subjected to the same treatment, namely NTERT1 and iPSCs exposed to simulated solar radiation (SSR), exhibited strong similarity (cosine similarity = 0.91), underscoring the consistent mutational response across disparate cellular backgrounds. Moreover, consistent with the colibactin-treated samples (**Figure 2E**), *SigRescueR* reliably produces highly stable signatures across replicates despite variability in exposure levels and baseline mutation burden (**Figures 3B, Supplementary Figure 3A**).

**Figure 3.**
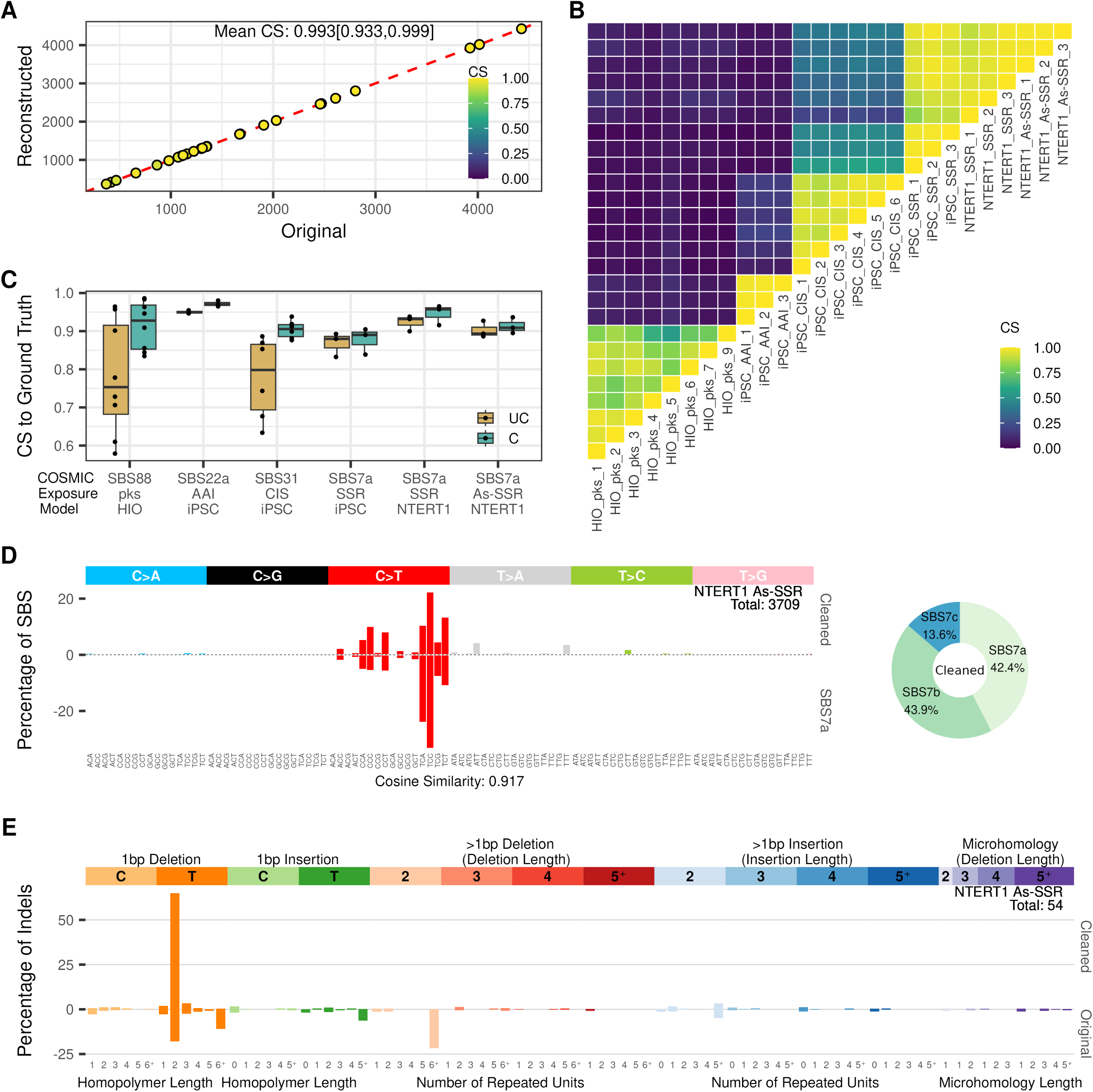
Performance evaluation of reconstruction, stability, and fidelity compared to the original and ground truth profiles. **(A)** Scatter plot comparing total mutation burden between original and reconstructed mutational profiles. Each dot represents a single sample exposure experiment and is colored according to their cosine similarity score between the original and reconstructed profiles. The cosine similarity score is summarized by the mean, with minimum and maximum values indicated in brackets. **(B)** Heatmap showing pairwise cosine similarity between each model and its replicates. **(C)** Boxplot summarizing cosine similarity to their COSMIC reference, separated by original (UC = uncleaned) and inferred exposure mutational signature (C = cleaned). Each dot represents a single sample. **(D)** Left panel: SBS-96 mutational signature plot comparing the inferred exposure (Cleaned) to SBS7a for NTERT1 exposed to SSR. The cosine similarity score is indicated between the cleaned signature and SBS7a. Right panel: Donut plot showing the relative contributions of COSMIC mutational signatures attributed to the cleaned signature. Each color represents a distinct COSMIC signature with percentages indicated. **(E)** ID-83 mutational signature plot comparing cleaned and original (uncleaned) profiles for NTERT1 co-exposed to SSR and arsenic. The total number of mutations in the cleaned profile is shown within the plots.

Notably, in certain instances, exposures may be non-mutagenic, resulting in mutational profiles indistinguishable from background patterns. Consequently, it is critical that *SigRescueR* attributes all mutations to background, avoiding false positive assignment. To evaluate this capability, we analyzed mouse intestinal organoid samples subjected to a high-fat diet (HFD) versus a standard diet (SD), known to generate highly similar profiles (cosine similarity = 0.993) [53]. Applying *SigRescueR* to the HFD samples with SD samples as background, we observed that nearly all mutations were correctly attributed to the background (**Supplementary Figure 3B**), with an average inferred exposure activity of 1.3%

Furthermore, a critical question emerged: do these inferred exposure mutational signatures capture biologically meaningful signatures or are they statistical artifacts of Bayesian inference? To address this, we assessed the concordance between the modeled exposure mutational signatures and their COSMIC reference. Compared to the original profiles, we observed a marked increase in cosine similarity to the reference (ANOVA *p*-value = 0.002) (**Figure 3C**). Notably, the magnitude of improvement and the ability of the inferred exposure mutation signature to approach near-perfect concordance varied between replicates—an expected reflection of inherent sample-to-sample biological variability. In models exposed to SSR and aristolochic acid I (AAI), the original profiles exhibited strong concordance with SBS7a, a UV-associated signature [55], and SBS22a, AAI-attributed signature [21,56], respectively, with cosine similarity greater than 0.9. Baseline correction further improved similarity. Conversely, two HIO treated with colibactin initially showed low similarity to SBS88 [25,26,54], but baseline correction substantially elevated cosine similarity from 0.58 to 0.86 and 0.61 to 0.84 in replicates 4 and 5, corresponding to improvements of 47.8% and 38.5%, respectively (**Figure 3C**). Collectively, these results demonstrate *SigRescueR*’s robustness to extract biologically meaningful signatures across diverse experimental contexts, while maintaining insensitivity to activity magnitude.

Next, we analyzed NTERT1 cells exposed to SSR, focusing on similarity to SBS7a [55]. *SigRescueR* demonstrated reliable baseline correction, with a mean baseline activity of 0.104, confirming its ability to preserve biologically relevant signals (**Figure 3D, left**). Given that SSR induces a complex mixture of UV-associated signatures, decomposition of the corrected profile revealed SBS7a, SBS7b, and SBS7c [57] as major contributors, explaining the cosine similarity of 0.92 observed (**Figure 3D, right**).

### Effective noise detection and baseline correction beyond SBS

Beyond SBS, baseline correction was applied to insertion-deletion mutations (indels) in NTERT1 cells co-exposed to SSR and arsenic [52]. This resulted in an increase in cosine similarity to ID13 [10], a UV-associated signature, between the original and cleaned profile, from 0.592 to 0.974 (**Figure 3E**). Moreover, correction of doublet base substitutions (DBS) maintained high concordance with the UV-associated DBS1 signature [58], with cosine similarity reaching 0.994 (from 0.986 to 0.994; **Supplementary Figure 4**).

We further validated *SigRescueR*’s versatility using indel mutations from HIO exposed to colibactin. Consistent high cosine similarity was observed, increasing from 0.94 to 0.97 toward the colibactin-associated ID18 signature [54] (**Supplementary Figure 5A**). Although an established DBS signature for colibactin is lacking, applying baseline correction to DBS mutations revealed a baseline activity of 10.3%, indicative of effective noise detection (**Supplementary Figure 5B**). Collectively, these findings highlight *SigRescueR*’s robustness and broad applicability across various mutational categories.

### Accuracy of baseline correction using strand-aware SBS classification

Accurate identification of transcription strand bias (TSB) is essential to understand how specific agents cause strand-specific DNA damage and repair, revealing key drug mechanisms and mutational outcomes [10,59]. To further elucidate the impact of (TSB) on mutational signature rescue, we extended our analysis to SBS288 classifications. We applied *SigRescueR* across a range of biologically relevant samples: MEFs exposed to benzo[a]pyrene (B[a]P), AAI and ultraviolet light (UVC); BEAS-2B exposed to B[a]P; iPSCs exposed to AAI, cisplatin, and SSR; and colibactin-treated HIO [21,25,34,51,52] (**Figure 4**, **Supplementary Figure 5C**).

**Figure 4.**
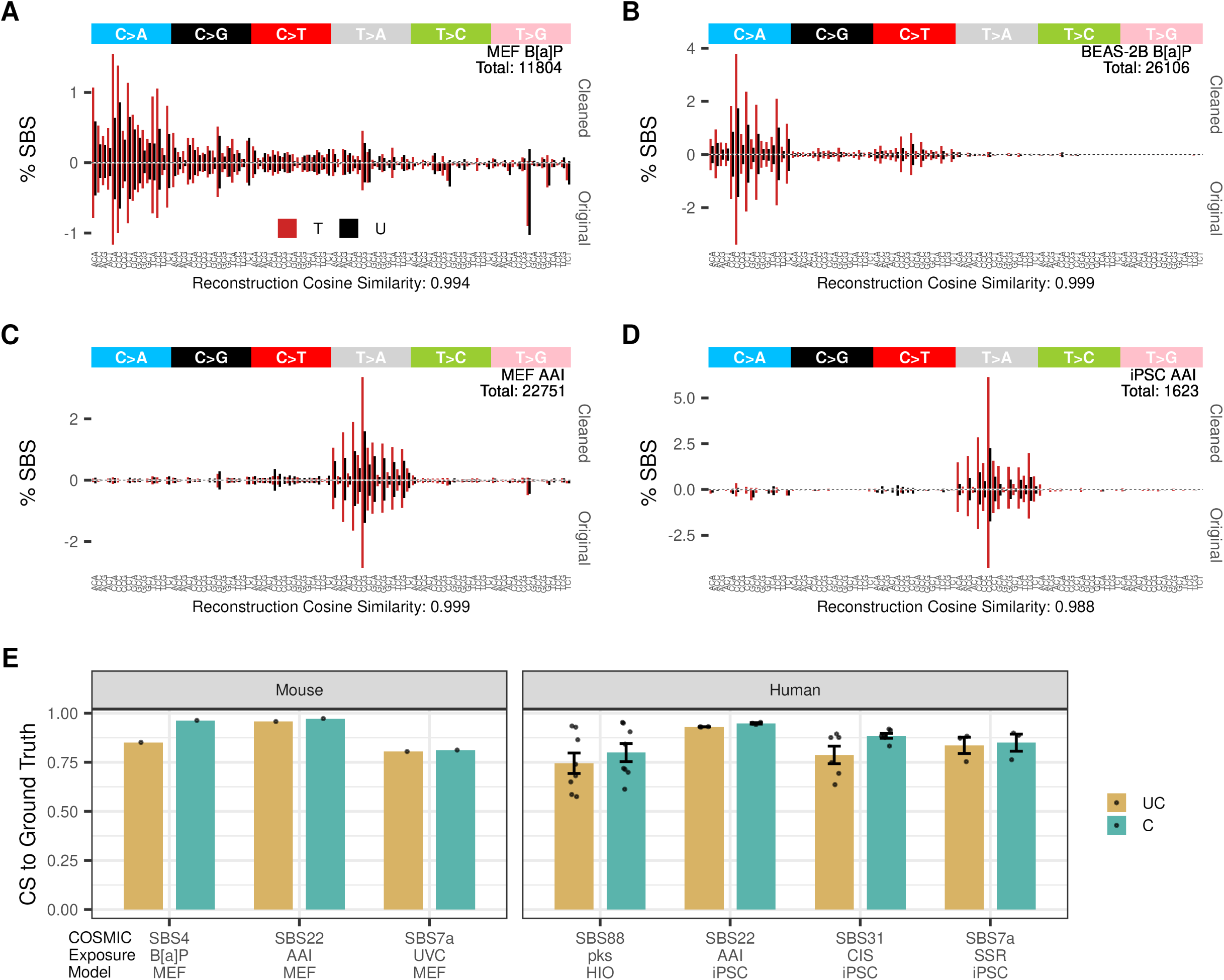
Beyond SBS96: integrating transcription strand bias Information for cleanup. SBS-288 mutational signature plots show cleaned and original mutational spectra for **(A)** MEF exposed to B[a]P, **(B)** BEAS-2B exposed to B[a]P, **(C)** MEF exposed to AAI, and **(D)** iPSC exposed to AAI. Colors indicate mutations on transcribed (T) and untranscribed (U) strands. **(E)** Barplot summarizing cosine similarity to the reference, separated by original (UC = uncleaned) and inferred exposure mutational signatures (C = cleaned). Each dot represents a single sample. The total mutation count in the cleaned profiles is indicated within the plots. Cosine similarity scores compare reconstructed and original mutational spectra.

Incorporating TSB information reinforced the robustness of our approach across more granular mutational classifications. Similarity to the tobacco-associated signature SBS4 [5] improved in MEFs, rising from 0.85 to 0.958 at the SBS96 level (**Supplementary Figure 6A**) and further to 0.963 at the SBS288 level (**Figure 4A**), while exposure activity remained stable (**Supplementary Figure 6B**). In BEAS-2B cells exposed to B[a]P, the cosine similarity before and after cleanup using the SBS288 channel remained stable, with approximately 6% baseline profile removed, while preserving TSB (**Figure 4B**). Parallel improvements were observed in MEFs treated with AAI and UVC when compared to SBS22a [21,56] and SBS7a [55], respectively (**Figure 4C**, **Supplementary Figure 6C&D**). Similarly, in AAI-treated iPSCs, cosine similarity increased modestly from 0.939 to 0.953 (**Figure 4D**). Overall, *SigRescueR* maintains or improves cosine similarity with strand bias information (**Figure 4E**) without affecting reconstruction accuracy (average cosine similarity of 0.98) (**Figure 4A-D**), demonstrating that *SigRescueR*’s accuracy extends well to strand-aware mutational signature classifications.

### Benchmarking against existing methods demonstrates rigor and outperformance

We benchmarked *SigRescueR* against several established approaches, including simple subtraction [25,34], NNLS [39], and *SparseSignatures [38]*. Given NNLS’s known propensity for overfitting [40], we implemented a 90% shrinkage strategy, where only 90% of the inferred baseline mutational signature is subtracted from the observed profile.

The main premise of baseline correction through signature decomposition is to parse the mutational spectra into known and, where applicable, latent signatures, while preserving the total mutation count. Across the four evaluated methods, *SigRescueR* excelled by closely matching the total mutation burden between the reconstructed and the original profiles, outperforming simple subtraction (53.3 [32.5; 74.1]), NNLS (561 [236,886]), and *SparseSignatures* (-77.1 [-95.8; -53.8]) (**Figure 5A**). Notably, subtraction and NNLS often attributed baseline activity per mutation context that exceeded the original mutation counts, resulting in negative mutation contexts, averaging 26 and 51 negative channels, respectively. Since negative mutations are biologically implausible and indicative of modeling artifacts, truncating to zero elevated mutation counts during reconstruction. Conversely, *SparseSignatures* underestimated mutation counts by leaving some mutations unassigned, resulting in lower reconstructed mutations. This comparison underscores *SigRescueR*’s balanced ability to accurately reconstruct mutation counts while avoiding biologically implausible artifacts. While *SparseSignatures* demonstrated remarkable alignment between reconstructed and original spectra, *SigRescueR* also delivered excellent performance, achieving a mean cosine similarity of 0.981 across 26 tested samples (**Figure 5B**).

**Figure 5.**
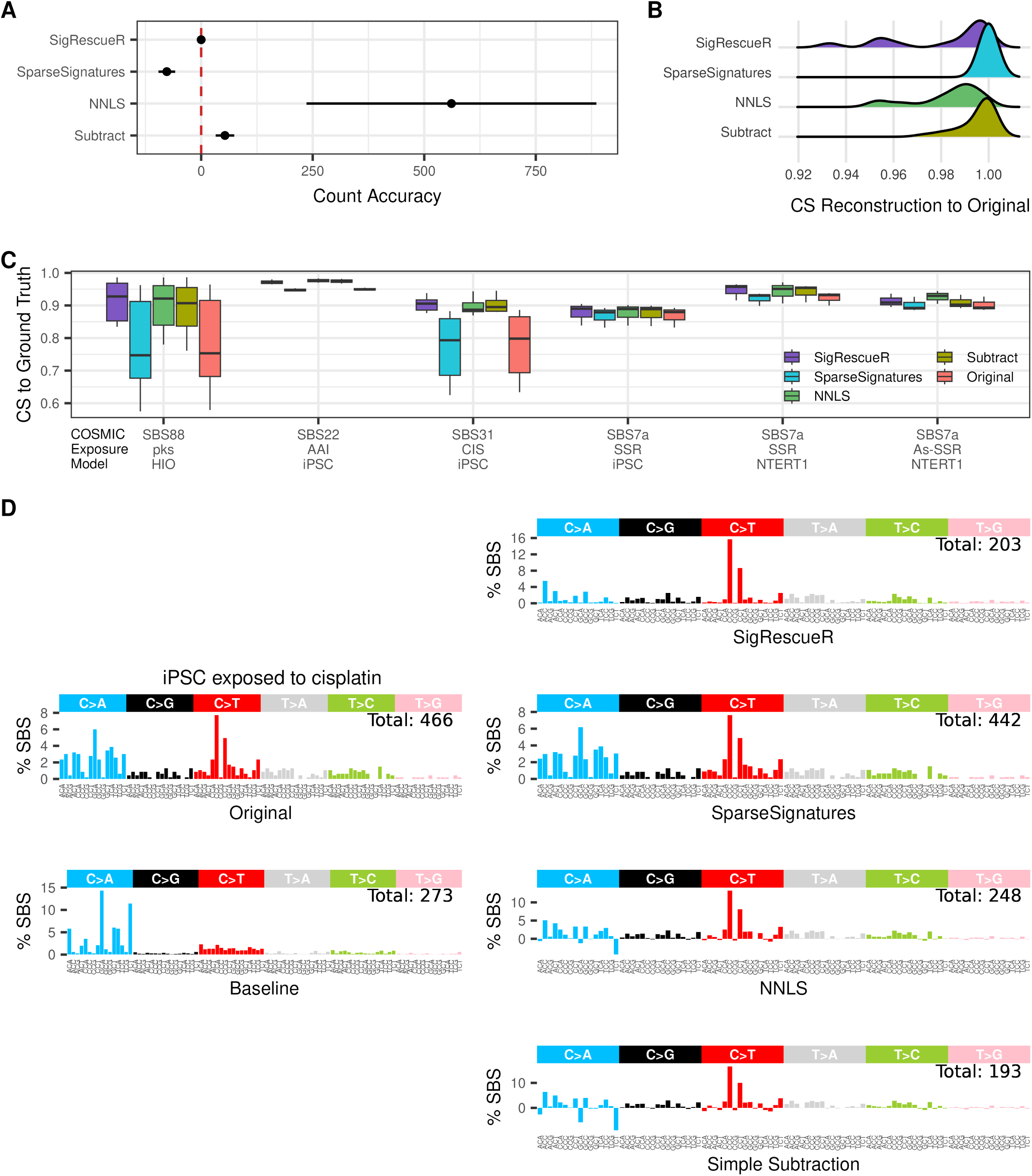
Model comparison of reconstruction accuracy, stability, and fidelity. **(A)** Dot-and-whisker plot depicting count accuracy measured as the difference in total mutation count between reconstructed and observed profiles. Each dot represents the mean, and whiskers indicate the 95% range across all samples. The red dashed line marks perfect count accuracy. **(B)** Distribution plot of cosine similarity between reconstructed and observed mutational spectra**.(C)**Boxplot summarizing cosine similarity to the reference, separated by cleaning methods. **(D)** SBS-96 mutational signature plots showing the uncleaned (Original), untreated (Baseline), and the inferred exposure mutational signatures by four methods using iPSC exposed to cisplatin. Total mutation counts in cleaned profiles are indicated within the plots.

Moreover, we computed cosine similarity for exposure signatures inferred from simple subtraction, NNLS and *SparseSignatures,* against COSMIC reference, using *SigRescueR* as the benchmark (**Figure 5C)**. *SigRescueR* consistently demonstrated superior performance over competing methods, with variations in cosine similarity among top-performing samples averaging only 0.0125 (**Figure 5C**). **Figure 5D** illustrates the signatures inferred across the four methods for cisplatin-treated iPSCs. Simple subtraction and NNLS yielded similar cosine similarity scores to SBS31 [24], a platinum-based chemotherapy COSMIC signature, at 0.88 and 0.87, respectively, achieved by clipping negative channels to zero. In contrast, *SparseSignatures* failed to detect any baseline signatures, while *SigRescueR* effectively removed baseline mutations and achieved the highest cosine similarity score equaling 0.903 to SBS31 signature.

### Case studies using simulated and real patients’ tumor data

Unlike controlled experimental models, the human cancer genome accumulates mutations from a complex interplay of multiple mutational processes [10,11], making it inherently noisier. We evaluated *SigRescueR*’s effectiveness in detecting and removing artifactual signatures, specifically SBS45 [32] and SBS54 [10], by injecting varying levels of each signature into 100 randomly selected negative samples as confirmed by SigProfilerAssignment [48] (**Supplementary Figure 7A**). Further, we applied the same evaluation to NNLS [46] and *SparseSignatures* [47] for benchmarking. To assess performance, we computed precision and sensitivity (**Figure 6, Supplementary Figure 7A**). Among the three evaluated tools, *SigRescueR* achieved the highest overall F1 score for both SBS45 and SBS54, followed by NNLS and then *SparseSignatures* (**Figure 6**). While *SigRescueR* and NNLS both displayed high sensitivity, *SigRescueR* outperformed NNLS by producing fewer false-positive calls, 0.956 and 0.879 for SBS45, and 0.883 and 0.859 for SBS54, respectively. In contrast, *SparseSignatures* showed inconsistent performance, with high F1 scores for some samples and failing to detect the signatures in others (**Supplementary Figure 7A**). Interestingly, several samples injected with SBS45 resulted in a precision less than 0.9 (n = 11), with precision dropping as low as 0.576 in one sample. Of the 11 samples, the true level of injections fell within the credible interval for six samples, appropriately accounting for the uncertainty. For the five remaining samples, four samples lied slightly outside the credible interval with a difference of 0.01 from the true injection value, while the fifth sample had a difference of 0.14. The overcalled mutations likely are a result of SBS45-like patterns within other COSMIC signatures. Applying *SigRescueR* across all COSMIC signatures revealed SBS45 attributions (**Supplementary Table 4**). For example, when analyzing SBS4, approximately 33% of extracted mutations were attributed to SBS45. Thereafter, we reassessed these five samples by combining the SBS45 attribution detected before injection with the true SBS45 injection values. This operation accounted for the undetected SBS45, resulting in an estimate that aligns with the initial posterior distribution and confirming that the low precision was due to undetected SBS45 signatures present in the original samples. For example, in PD51293a, the level of injection was 21%, yet the credible interval spanned 22-27%. Cleaning the original profile revealed an additional 4.4% SBS45, which brings the estimate within the credible interval. Moreover, when comparing the original to the cleaned injected spectra, we observed strong preservation of the original spectra with a mean cosine similarity of 0.97 and 0.98 for SBS45 and SBS54, respectively (**Supplementary Figure 7B**).

**Figure 6.**
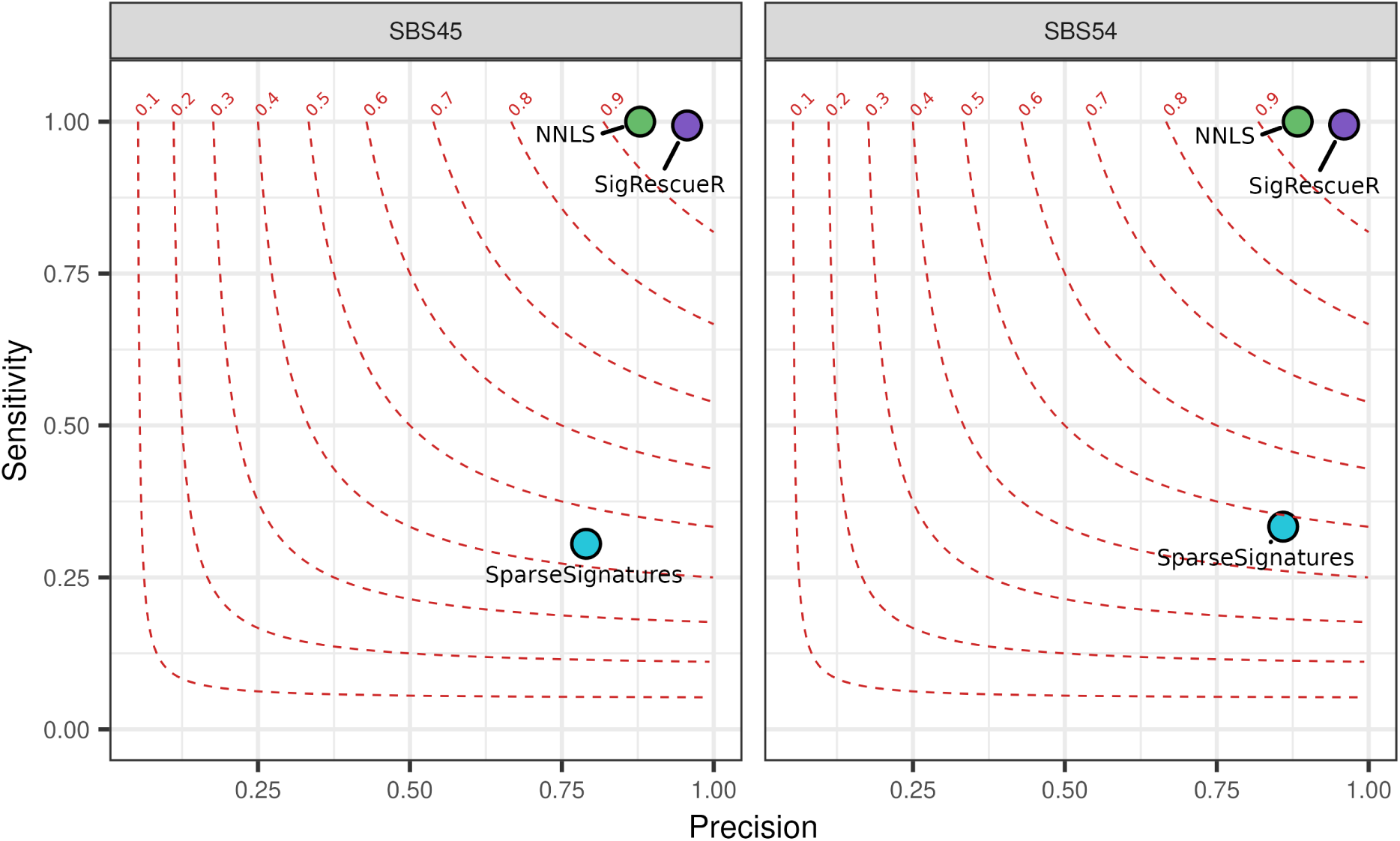
*SigRescueR* performance on simulated and real tumor data. Scatter plot showing the average precision and sensitivity for cleaning SBS45 or SBS54 across 100 randomly selected samples negative for these signatures. Results are compared between *SigRescueR*, *SparseSignatures*, and NNLS. Each dot is colored according to the cleaning method used.

After validating *SigRescueR*’s performance, we applied the tool to real tumor samples harboring SBS45 and SBS54 signatures. *SigRescueR* effectively removed these signatures, as confirmed by SigProfilerAssignment [48] (**Supplementary Figure 7C-D**). Importantly, we found that cleaning SBS45 preserved the integrity of existing SBS signatures. However, in one sample, it led to reassignment between two signatures with overlapping profiles. Specifically, SBS41 was reassigned to SBS93, and SBS5 to SBS40a (**Supplementary Figure 7D**). This finding demonstrates *SigRescueR*’s high sensitivity in accurately distinguishing overlapping mutational patterns without broadly disrupting the mutational landscape. Furthermore, by removing artifactual signatures, *bona fide* signatures can emerge more clearly, offering biologically meaningful insights into the mutational processes driving cancer development. This underscores *SigRescueR*’s value not only for refining mutational profiles but also as a powerful tool for uncovering clinically relevant signatures.

### Extending *SigRescueR* applications to duplex sequencing data

Error-corrected next-generation sequencing (ecNGS) is an emerging technology that provides substantial advantages over conventional whole-genome sequencing approaches by markedly reducing sequencing errors and enhancing the detection of low-frequency variants [13,60]. Among ecNGS platforms, single-molecule duplex sequencing, based on sequencing both DNA strands independently to build consensus variant calls, has gained prominence as one of the most innovative and accurate methods available today [60]. Several duplex-based frameworks have been developed to improve the detection of exposure-related mutational signatures without the need for clonal expansion, thereby minimizing culture-induced artifacts [61–64] . However, despite their precision, these methods remain prone to technology-related artifacts. These residual artifacts obscure true biological signals and complicate the identification of true mutational signatures.

To address this challenge, we applied *SigRescueR* on duplex sequencing data from patients treated with 5-fluorouracil (5-FU) [65] using naïve and memory B and T cells, and monocytes. *SigRescueR* enabled the recovery of SBS17b, a COSMIC signature commonly ascribed to 5-FU treatment [66,67] (**Figure 7A&B**). In one patient, attribution to SBS17b increased from 411 to 571. Notably, *SigRescueR* revealed attribution to SBS17b in eight 5-FU samples that were previously negative for this signature (**Figure 7B**), highlighting *SigRescueR*’s capability to recover treatment-induced signatures that were previously undetected.

**Figure 7.**
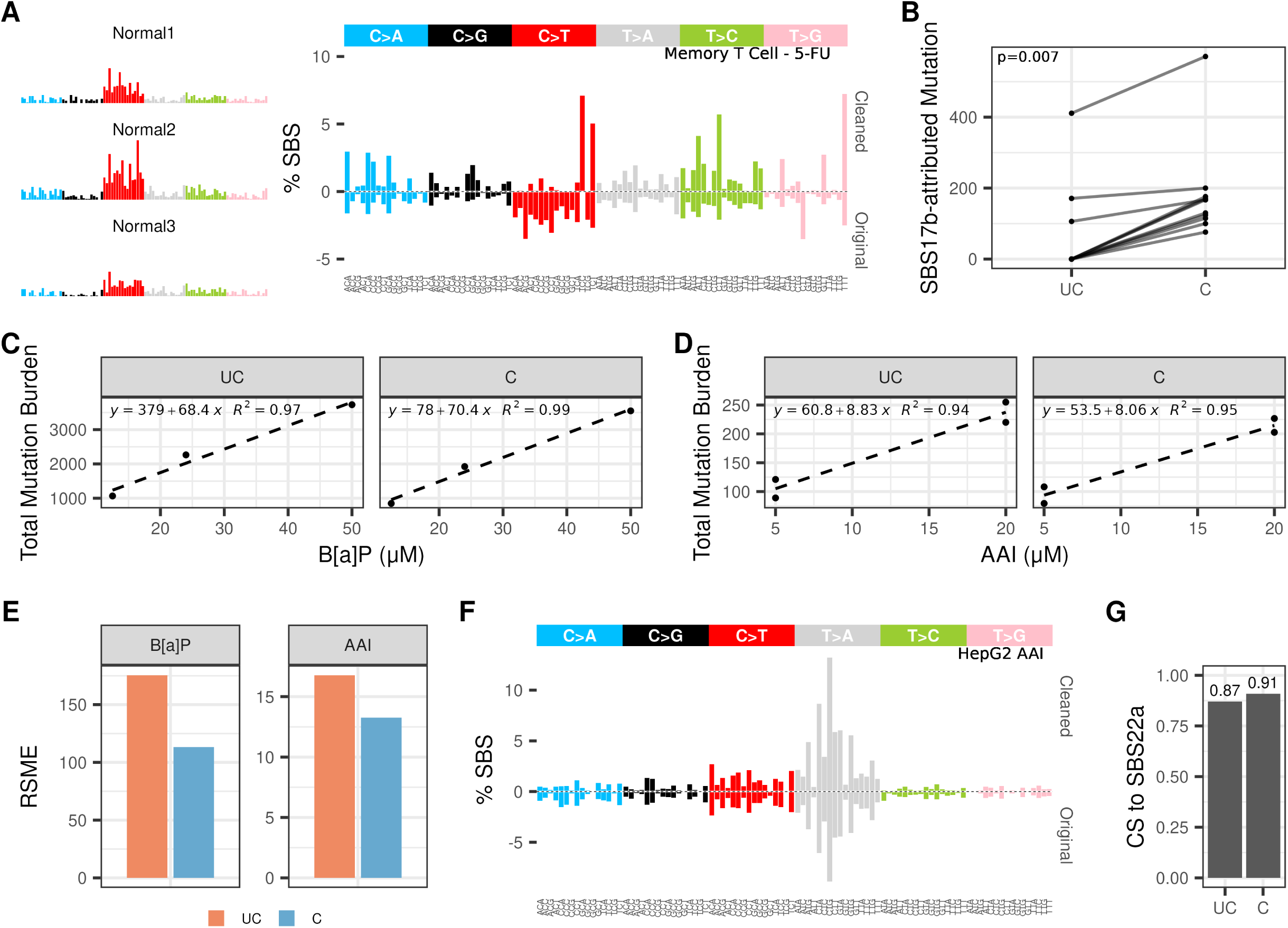
*SigRescueR* performance with ecNGS. **(A)** SBS-96 mutational signature plots showing three unexposed memory T cell profiles (Normal) and inferred exposure signatures (Cleaned) alongside uncorrected original mutational spectra (Original) from an individual previously exposed to 5-fluorouracil. **(B)** Connected dot plot illustrating mutation counts attributed to SBS17b before (UC = uncleaned) and after cleaning (C = cleaned) across various individuals and cell types; each connected pair represents one sample. Statistical significance (*p*-value) was calculated using the Wilcoxon Rank Sum Test. Scatter plots showing mutation counts for **(C)** B[a]P and **(D)** AAI across different concentrations before and after cleaning; each dot is a sample; fitted linear regression lines with equation and R^2^ values are shown. **(E)** Barplot showing residual error measured by root mean squared error (RSME), separated by original (UC = uncleaned) and cleaned (C = cleaned) profiles for B[a]P and AAI. **(F)** SBS-96 mutational signature plots comparing inferred exposure (Cleaned) and uncorrected spectra (Original) for the 3D HepG2 model exposed to AAI. **(G)** Barplot summarizing cosine similarity of the 3D HepG2 model to SBS22a between original (UC = uncleaned) and cleaned (C = cleaned) profiles.

Next, we utilized duplex sequencing data from a mouse bone marrow model exposed to B[a]P across different dosage levels [68]. *SigRescueR* maintained a clear dose-response relationship in mutational impact and enhanced the linear correlation between dose and mutation frequency, reflected by a 2% increase R² value (**Figure 7C**). Applying *SigRescueR* on 3D HepG2 spheroids exposed to AAI [69] revealed a congruent trend, where R^2^ increased from 0.94 to 0.95 (**Figure 7D**). This improvement in linear relationship also resulted in a reduction in the root mean squared error (RMSE) (**Figure 7E**), indicating less deviation and variability between replicates, which is crucial for downstream analyses in cancer genomics and toxicology studies [13]. Furthermore, *SigRescueR* improved the cosine similarity to SBS22a from 0.87 to 0.91, validating its ability to refine ecNGS-derived mutational landscapes (**Figure 7F&G**).

Collectively, these results demonstrate that *SigRescueR* not only removes low-confidence artifacts and sequencing noise from raw variant calls but also enhances the detection and interpretability of true mutational signatures. This approach establishes a high-confidence framework for ecNGS-based mutagenesis studies, expanding the analytical capabilities of duplex sequencing technologies.

## Discussion

Deconvolution of mutational signatures is fundamental to understanding the complex mutational processes underlying cancer and environmental mutagenesis. However, accurately distinguishing true exposure-related signatures from background noise remains a persistent challenge due to sequencing errors and biological variability [7,10,28,32,33]. Our study demonstrates that *SigRescueR* addresses this challenge by implementing a robust baseline correction approach, effectively revealing *bona fide* mutational signals that are otherwise obscured by background noise.

*SigRescueR* is designed around three core strategies. First, it applies a strong baseline correction, carefully avoiding negative values, ensuring the inferred exposure mutation signature remains biologically meaningful. Second, it considers the overall shape of the observed by assessing how well the observed data fit the reconstructed profile using a Dirichlet distribution. Third, a COM-Poisson likelihood [41], penalizes large deviations from the observed count, preventing the exposure profile from absorbing leftover counts. The tool is well-suited for single-sample analyses.

To evaluate *SigRescueR*’s performance, we applied the tool across a broad spectrum of experimental models, incorporating murine and human systems with well-characterized spectra, in addition to patient-derived tumor datasets. We demonstrate that *SigRescueR* not only enhances the fidelity of reconstruction, but also reliably distinguishes true exposure signatures from non-mutagenic backgrounds, limiting false positive assignments, a critical feature for high-confidence mutational inference. Moreover, *SigRescueR*’s shows high reproducibility across biological replicates, even under variable exposure intensities, providing a key advantage for mutagenesis studies and mechanistic toxicology[14].

Moreover, we showcased *SigRescueR*’s ability to deliver consistent results across SBS96 and SBS288 classifications, as well as for indels and DBS. Compared to existing methods [25,34,46,47], *SigRescueR* consistently matches or surpasses their performance, demonstrating superior accuracy and robustness. We show that simple subtraction [25,34] and NNLS [46] can generate negative values when the background burden is higher than the exposure. In contrast, *SparseSignatures* achieves high reconstruction accuracy [47], but this can cause background mutations being absorbed by the exposure signature.

Furthermore, we challenged *SigRescueR* to refine tumor sequencing data by removing sequencing-related artifacts. This task is particularly difficult due to the absence of a clearly defined true background and the complex mixture of mutational processes accumulated over time in tumor samples [5,15,16]. Our results demonstrate that *SigRescueR* effectively removes sequencing artifacts with high sensitivity and precision. Importantly, *SigRescueR* can be applied flexibly to clean other signatures beyond sequencing noise, such as SBS17, which is relevant when analyzing mouse samples [29], as well as APOBEC-related signatures (SBS2 and SBS13) [15,22] that often dominate mutational spectra in patients. This adaptability addresses cases where signature attribution may be ambiguous, with some signatures potentially unassigned and absorbed by others, a challenge *SigRescueR* can identify and manage [70].

Leveraging duplex sequencing data, *SigRescueR*’s framework can detect novel low-frequency mutational signatures often obscured by technical artifacts, while also reducing variability between replicates. This capability expands our understanding of mutational heterogeneity, uncovers previously unrecognized mutational mechanisms contributing to carcinogenesis, and supports toxicology and cancer etiology studies.

Despite *SigRescueR*’s overall strength, we note several limitations. First, *SigRescueR* relies on Bayesian inference, producing posterior distribution rather than point estimates, that are summarized as posterior mean. We treat the posterior mean as point estimates, despite it being the most probable value rather than the value that minimizes the residual error. Second, *SigRescueR* is agnostic to the level of background, bounded by the priors, and will remove as much as the posterior determines given the data. If the default settings for summarizing the posterior does not meet the user’s needs, we provide the flexibility to explore the posterior distribution. Third, the computational time is directly proportional to mutation burden as *SigRescueR* computes the normalizing constant □ for all mutation context required for the COM-poisson likelihood to ensure the probabilities sum up to one. Despite these limitations, *SigRescueR* provides a full distribution that can be used to assess uncertainty through credible intervals and confidence levels, rather than committing to a single estimate that likely reflects a minimization.

Taken altogether, our results demonstrate that *SigRescueR* enhances the accuracy of mutational signature analysis by rescuing biologically meaningful signals obscured by background noise, sequencing artifacts, or modeling bias, enabling new biological discoveries in data that would otherwise remain obscured. It consistently outperforms existing methods in reconstructing mutational signatures with near-perfect fidelity. Its versatility across diverse experimental models, species, sequencing technologies, and mutation classifications highlights its broad applicability. *SigRescueR* is a first-of-its-kind tool able to reliably identify true mutational signatures from exposure models, even with limited sample numbers. These qualities position it as a powerful tool for mutational signature analysis, capable of advancing studies in cancer genomics and other fields reliant on precise mutation pattern characterization. Future work can further extend its utility to additional data types, experimental conditions, and sequencing platforms fostering deeper insights into mutagenic processes.

## Code and data availability

*SigRescueR* is implemented in R and provided as open-source software on GitHub at https://github.com/ZhivaguiLab/SigRescueR/. All code used to generate the figures and analyses reported in this manuscript is openly available on GitHub at: https://github.com/ZhivaguiLab/SigRescueRmanu/. Detailed instructions for reproducing the results are provided in the repository’s README file.

## References

1. Hanahan D, Weinberg RA. The hallmarks of cancer. Cell 2000; 100:57–70

2. Hanahan D, Weinberg RA. Hallmarks of cancer: the next generation. Cell 2011; 144:646–674

3. Hanahan D. Hallmarks of cancer: New dimensions. Cancer Discov. 2022; 12:31–46

4. Vogelstein B, Papadopoulos N, Velculescu VE, et al. Cancer genome landscapes. Science 2013; 339:1546–1558

5. Alexandrov LB, Nik-Zainal S, Wedge DC, et al. Signatures of mutational processes in human cancer. Nature 2013; 500:415–421

6. Volkova NV, Meier B, González-Huici V, et al. Mutational signatures are jointly shaped by DNA damage and repair. Nat. Commun. 2020; 11:2169

7. Zou X, Koh GCC, Nanda AS, et al. A systematic CRISPR screen defines mutational mechanisms underpinning signatures caused by replication errors and endogenous DNA damage. Nat. Cancer 2021; 2:643–657

8. Otlu B, Alexandrov LB. Evaluating topography of mutational signatures with SigProfilerTopography. Genome Biol. 2025; 26:134

9. Alexandrov LB, Zhivagui M. Mutational signatures and the etiology of human cancers. Reference Module in Biomedical Sciences 2018;

10. Alexandrov LB, Kim J, Haradhvala NJ, et al. The repertoire of mutational signatures in human cancer. Nature 2020; 578:94–101

11. Helleday T, Eshtad S, Nik-Zainal S. Mechanisms underlying mutational signatures in human cancers. Nat. Rev. Genet. 2014; 15:585–598

12. Pfeifer GP. Environmental exposures and mutational patterns of cancer genomes. Genome Med. 2010; 2:54

13. Zhivagui M, Zavadil J. Mutational signatures in cancer genomics and toxicology. Comprehensive Toxicology 2026; 82–105

14. Zhivagui M, Korenjak M, Zavadil J. Modelling mutation spectra of human carcinogens using experimental systems. Basic Clin. Pharmacol. Toxicol. 2017; 121 Suppl 3:16–22

15. Nik-Zainal S, Alexandrov LB, Wedge DC, et al. Mutational processes molding the genomes of 21 breast cancers. Cell 2012; 149:979–993

16. Alexandrov LB, Nik-Zainal S, Wedge DC, et al. Deciphering signatures of mutational processes operative in human cancer. Cell Rep. 2013; 3:246–259

17. Alexandrov LB, Stratton MR. Mutational signatures: the patterns of somatic mutations hidden in cancer genomes. Curr. Opin. Genet. Dev. 2014; 24:52–60

18. Alexandrov LB, Jones PH, Wedge DC, et al. Clock-like mutational processes in human somatic cells. Nat. Genet. 2015; 47:1402–1407

19. Alexandrov LB. Understanding the origins of human cancer. Science 2015; 350:1175

20. Alexandrov LB, Zhivagui M. Mutational Signatures and the Etiology of Human Cancers. Encyclopedia of Cancer. 499–510

21. Nik-Zainal S, Kucab JE, Morganella S, et al. The genome as a record of environmental exposure. Mutagenesis 2015; 30:763–770

22. Chan K, Roberts SA, Klimczak LJ, et al. An APOBEC3A hypermutation signature is distinguishable from the signature of background mutagenesis by APOBEC3B in human cancers. Nat. Genet. 2015; 47:1067–1072

23. Chawanthayatham S, Valentine CC 3rd, Fedeles BI, et al. Mutational spectra of aflatoxin B1 in vivo establish biomarkers of exposure for human hepatocellular carcinoma. Proc. Natl. Acad. Sci. U. S. A. 2017; 114:E3101–E3109

24. Boot A, Huang MN, Ng AWT, et al. In-depth characterization of the cisplatin mutational signature in human cell lines and in esophageal and liver tumors. Genome Res. 2018; 28:654–665

25. Pleguezuelos-Manzano C, Puschhof J, Rosendahl Huber A, et al. Mutational signature in colorectal cancer caused by genotoxic pks+ E. coli. Nature 2020; 580:269–273

26. Boot A, Ng AWT, Chong FT, et al. Characterization of colibactin-associated mutational signature in an Asian oral squamous cell carcinoma and in other mucosal tumor types. Genome Res. 2020; 30:803–813

27. Huang MN, Yu W, Teoh WW, et al. Genome-scale mutational signatures of aflatoxin in cells, mice, and human tumors. Genome Res. 2017; 27:1475–1486

28. Koh G, Degasperi A, Zou X, et al. Mutational signatures: emerging concepts, caveats and clinical applications. Nat. Rev. Cancer 2021; 21:619–637

29. Riva L, Pandiri AR, Li YR, et al. The mutational signature profile of known and suspected human carcinogens in mice. Nat. Genet. 2020; 52:1189–1197

30. Zhivagui M, Ng AWT, Ardin M, et al. Experimental and pan-cancer genome analyses reveal widespread contribution of acrylamide exposure to carcinogenesis in humans. Genome Res. 2019; 29:521–531

31. Ivanov D, Hwang T, Sitko LK, et al. Experimental systems for the analysis of mutational signatures: no ‘one-size-fits-all’ solution. Biochem. Soc. Trans. 2023; 51:1307–1317

32. Costello M, Pugh TJ, Fennell TJ, et al. Discovery and characterization of artifactual mutations in deep coverage targeted capture sequencing data due to oxidative DNA damage during sample preparation. Nucleic Acids Res. 2013; 41:e67

33. Islam SMA, Díaz-Gay M, Wu Y, et al. Uncovering novel mutational signatures by de novo extraction with SigProfilerExtractor. Cell Genom. 2022; 2:None

34. Mingard C, Battey JND, Takhaveev V, et al. Dissection of cancer mutational signatures with individual components of cigarette smoking. Chem. Res. Toxicol. 2023; 36:714–723

35. Lee DD, Seung HS. Learning the parts of objects by non-negative matrix factorization. Nature 1999; 401:788–791

36. Zhivagui M, Ardin M, Ng AWT, et al. Experimental analysis of exome-scale mutational signature of glycidamide, the reactive metabolite of acrylamide. bioRxiv 2018;

37. Lin X, Boutros PC. Optimization and expansion of non-negative matrix factorization. BMC Bioinformatics 2020; 21:7

38. Lal A, Liu K, Tibshirani R, et al. De novo mutational signature discovery in tumor genomes using SparseSignatures. PLoS Comput. Biol. 2021; 17:e1009119

39. Lawson CL. Solving least squares problems. 1995;

40. Slawski M, Hein M. Non-negative least squares for high-dimensional linear models: Consistency and sparse recovery without regularization. ejs 2013; 7:3004–3056

41. Conway RW, Maxwell WL. A queuing model with state dependent service rates. Journal of Industrial Engineering 1962;

42. R Core Team. R: A Language and Environment for Statistical Computing. R Foundation for Statistical Computing, Vienna, Austria. 2021;

43. Stan Development Team. RStan: the R interface to Stan. 2025;

44. Menéndez ML, Pardo JA, Pardo L, et al. The Jensen-Shannon divergence. J. Franklin Inst. 1997; 334:307–318

45. Wild F. lsa: Latent Semantic Analysis. CRAN: Contributed Packages 2005;

46. Mullen KM, van Stokkum IHM. Nnls: The Lawson-Hanson algorithm for non-negative least squares (NNLS). CRAN: Contributed Packages 2007;

47. Mella L, Lal A, Angaroni F, et al. SparseSignatures: An R package using LASSO-regularized non-negative matrix factorization to identify mutational signatures from human tumor samples. STAR Protoc. 2022; 3:101513

48. Díaz-Gay M, Vangara R, Barnes M, et al. Assigning mutational signatures to individual samples and individual somatic mutations with SigProfilerAssignment. Bioinformatics 2023; 39:btad756

49. Mann HB, Whitney DR. On a Test of Whether one of Two Random Variables is Stochastically Larger than the Other. Ann. Math. Stat. 1947; 18:50–60

50. Girden E. ANOVA: Repeated Measures. Sage University papers. Quantitative applications in the social sciences, Vol. 84. 1991; 77:

51. Kucab JE, Zou X, Morganella S, et al. A compendium of mutational signatures of environmental agents. Cell 2019; 177:821–836.e16

52. Speer RM, Nandi SP, Cooper KL, et al. Arsenic is a potent co-mutagen of ultraviolet light. Commun. Biol. 2023; 6:1273

53. Meyenberg M, Hakobyan A, Papac-Milicevic N, et al. Mutational landscape of intestinal crypt cells after long-term in vivo exposure to high fat diet. Sci. Rep. 2023; 13:13964

54. Lee-Six H, Olafsson S, Ellis P, et al. The landscape of somatic mutation in normal colorectal epithelial cells. Nature 2019; 574:532–537

55. Hayward NK, Wilmott JS, Waddell N, et al. Whole-genome landscapes of major melanoma subtypes. Nature 2017; 545:175–180

56. Senkin S, Moody S, Díaz-Gay M, et al. Geographic variation of mutagenic exposures in kidney cancer genomes. Nature 2024; 629:910–918

57. Saini N, Roberts SA, Klimczak LJ, et al. The impact of environmental and endogenous damage on somatic mutation load in human skin fibroblasts. PLoS Genet. 2016; 12:e1006385

58. Chen J-M, Férec C, Cooper DN. Patterns and mutational signatures of tandem base substitutions causing human inherited disease. Hum. Mutat. 2013; 34:1119–1130

59. Vöhringer H, Van Hoeck A, Cuppen E, et al. Learning mutational signatures and their multidimensional genomic properties with TensorSignatures. Nat. Commun. 2021; 12:3628

60. Marchetti F, Cardoso R, Chen CL, et al. Error-corrected next generation sequencing - Promises and challenges for genotoxicity and cancer risk assessment. Mutat. Res. Rev. Mutat. Res. 2023; 792:108466

61. Hoang ML, Kinde I, Tomasetti C, et al. Genome-wide quantification of rare somatic mutations in normal human tissues using massively parallel sequencing. Proc. Natl. Acad. Sci. U. S. A. 2016; 113:9846–9851

62. Abascal F, Harvey LMR, Mitchell E, et al. Somatic mutation landscapes at single-molecule resolution. Nature 2021; 593:405–410

63. Nandi SP, Cheng Y, Al-Azzam S, et al. A universal duplex sequencing approach for accurate detection of somatic mutations. bioRxivorg 2025;

64. Bae JH, Liu R, Roberts E, et al. Single duplex DNA sequencing with CODEC detects mutations with high sensitivity. Nat. Genet. 2023; 55:871–879

65. Mitchell E, Pham MH, Clay A, et al. The long-term effects of chemotherapy on normal blood cells. Nat. Genet. 2025; 57:1684–1694

66. Christensen S, Van der Roest B, Besselink N, et al. 5-Fluorouracil treatment induces characteristic T>G mutations in human cancer. Nat. Commun. 2019; 10:4571

67. Secrier M, Li X, de Silva N, et al. Mutational signatures in esophageal adenocarcinoma define etiologically distinct subgroups with therapeutic relevance. Nature Genetics 2016; 48:1131–1141

68. LeBlanc DPM, Meier M, Lo FY, et al. Duplex sequencing identifies genomic features that determine susceptibility to benzo(a)pyrene-induced in vivo mutations. BMC Genomics 2022; 23:542

69. Conway GE, Chavanel B, Virard F, et al. Harnessing the power of an advanced in vitro 3D liver model and error-corrected duplex sequencing for the detection of mutational signatures. Mutagenesis 2025;

70. Koh G, Zou X, Nik-Zainal S. Mutational signatures: experimental design and analytical framework. Genome Biol. 2020; 21:37

