## Supplemental figures for "*SigRescueR*: A Pan-System Framework for Noise Correction and Mutational Signature Identification Across Sequencing Platforms"

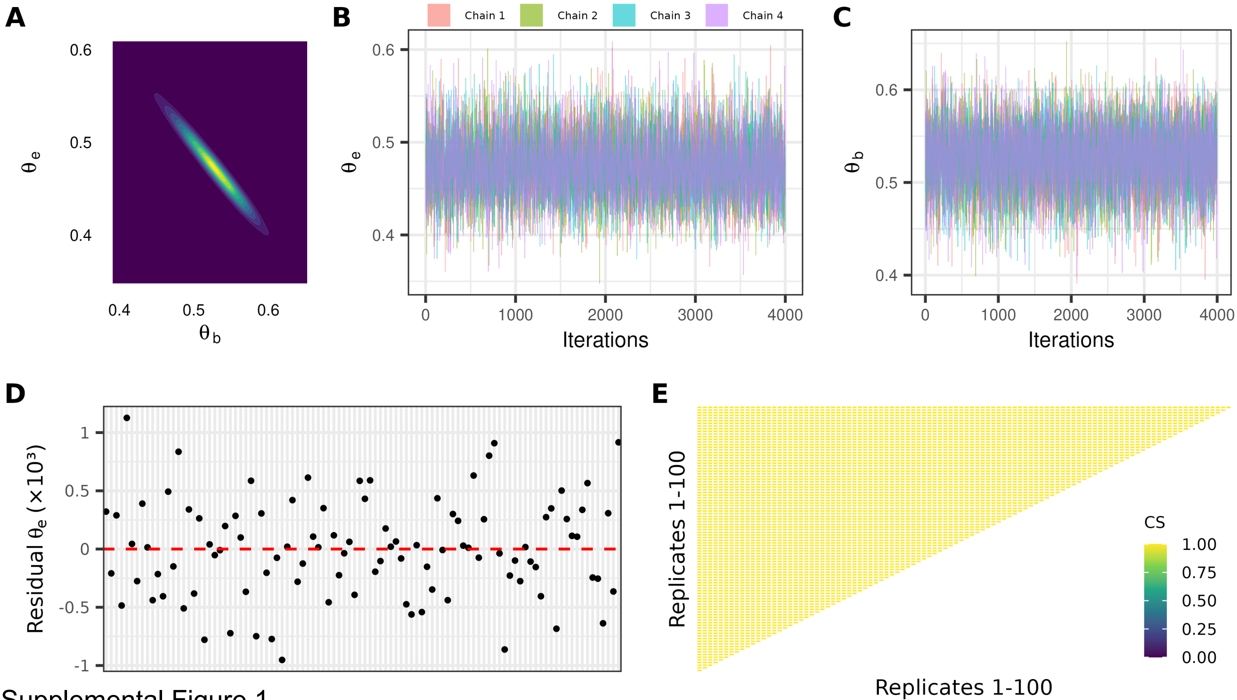


Supplemental Figure 1

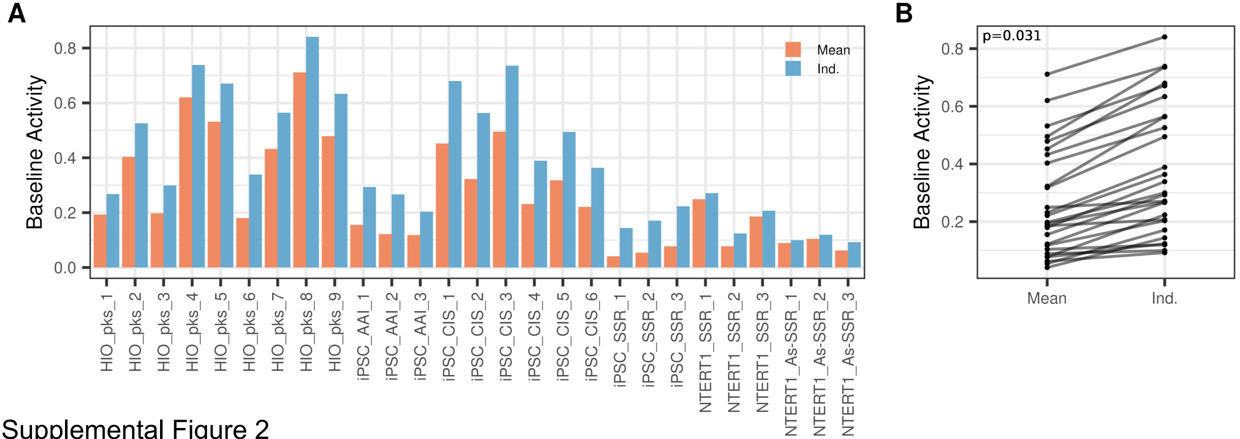


Supplemental Figure 2

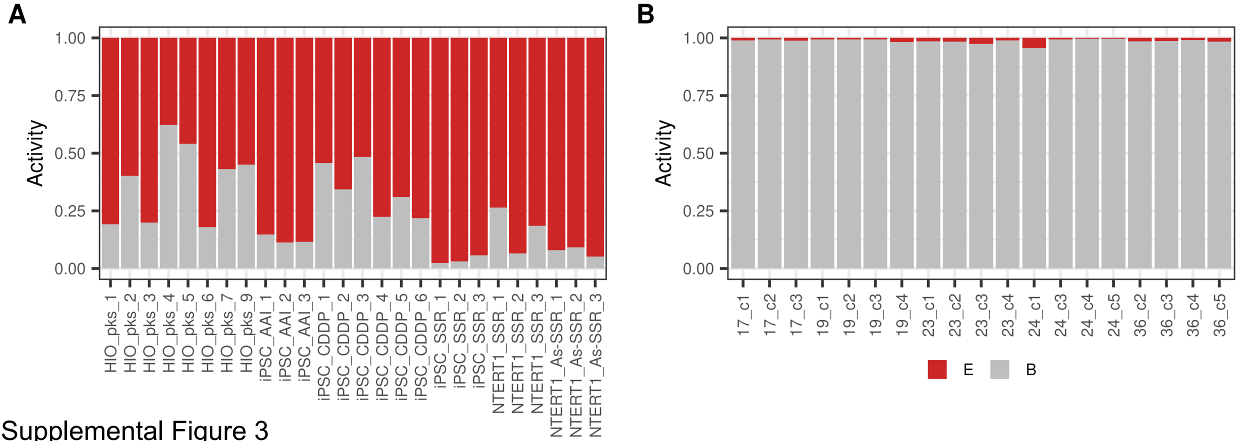


Supplemental Figure 3

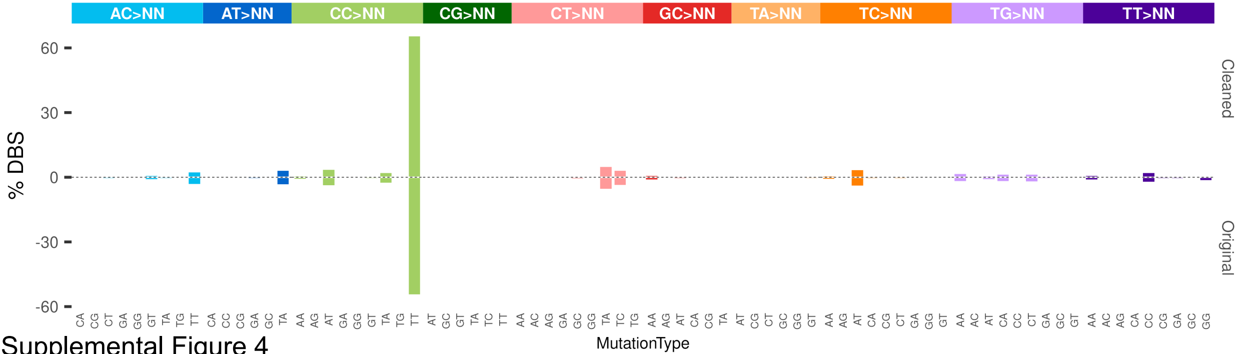


Supplemental Figure 4

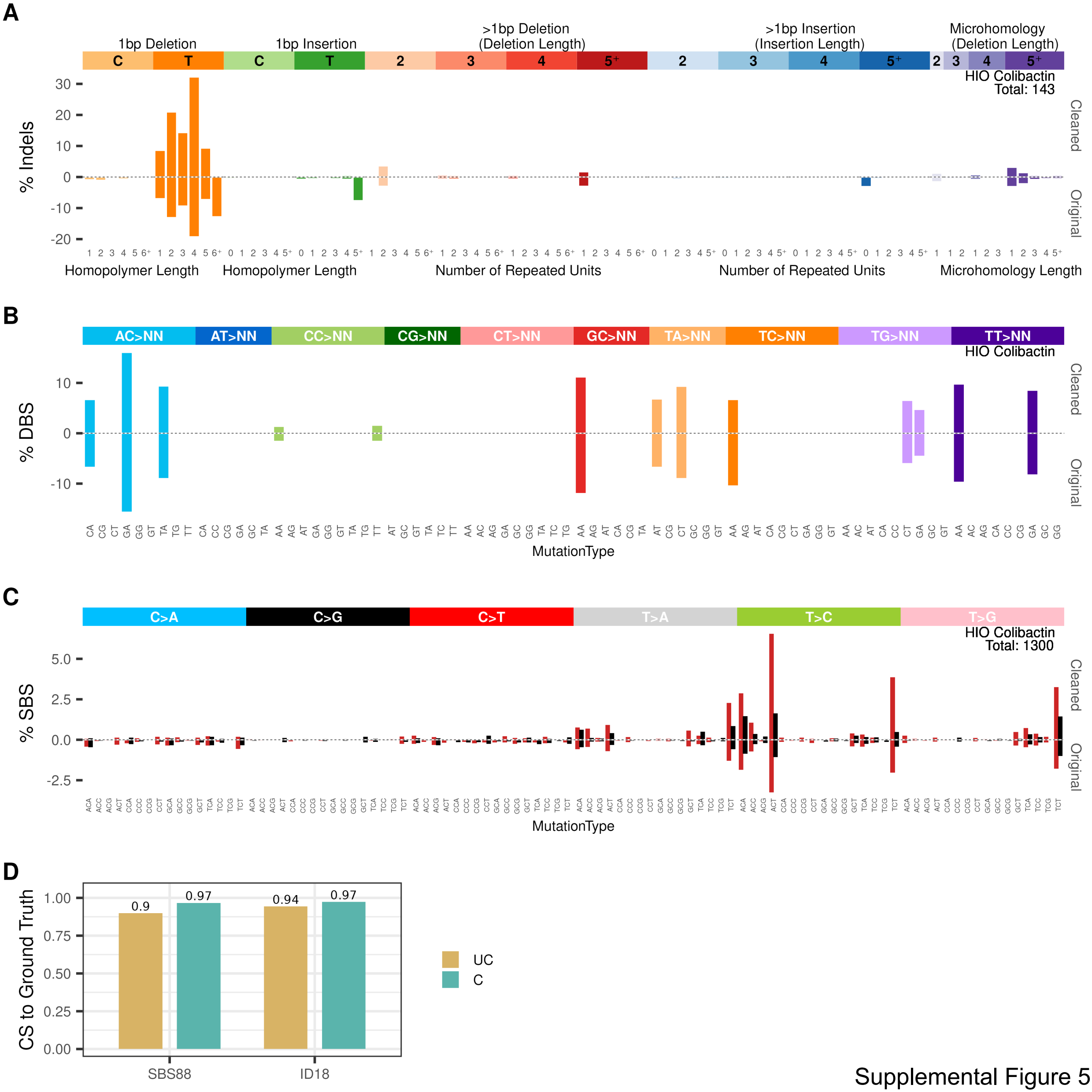


Supplemental Figure 5

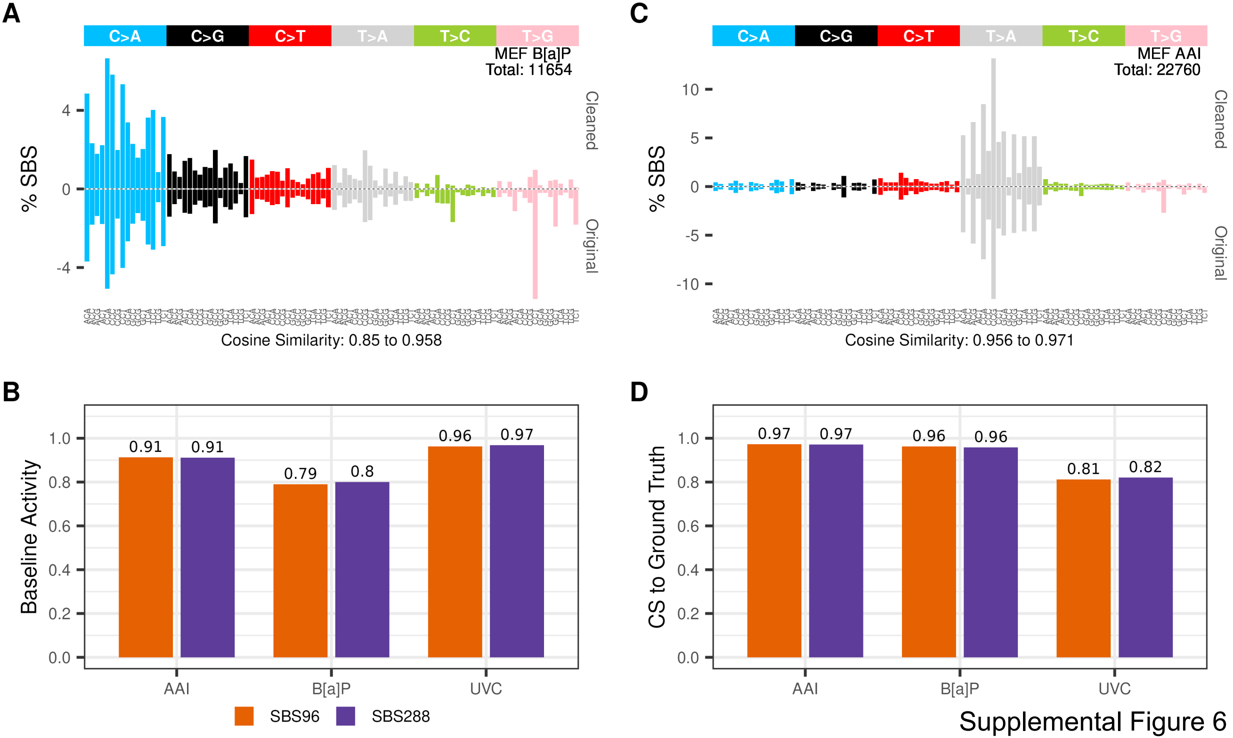


Supplemental Figure 6

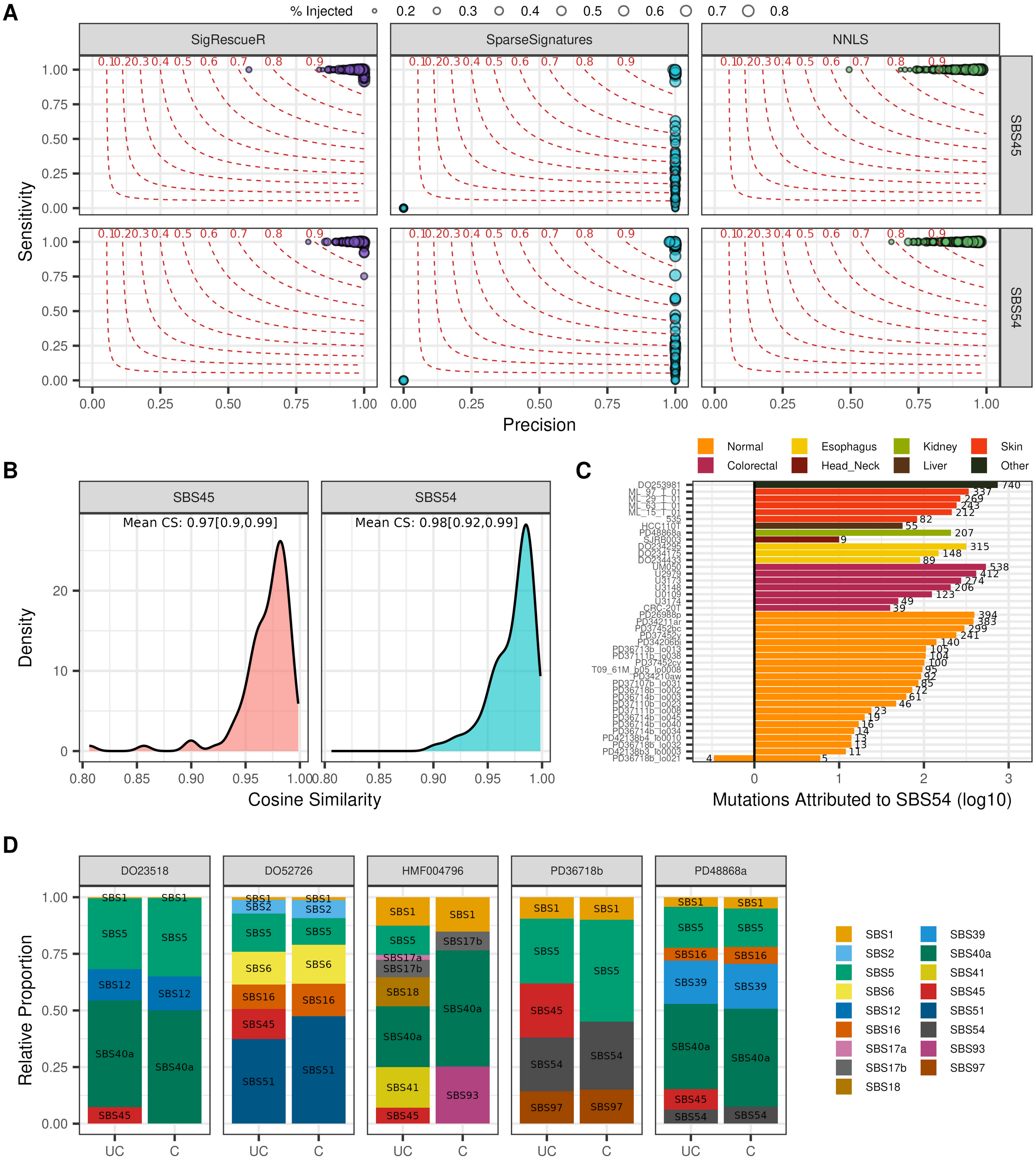


Supplemental Figure 7
